# Dynamics of human telomerase recruitment depend on template-telomere base-pairing

**DOI:** 10.1101/217885

**Authors:** Jens C. Schmidt, Arthur J. Zaug, Regina Kufer, Thomas R. Cech

## Abstract

The reverse transcriptase telomerase adds telomeric repeats to chromosome ends to counteract telomere shortening and thereby assures genomic stability in dividing human cells. Key variables in telomere homeostasis are the frequency with which telomerase engages the chromosome end and the number of telomeric repeats it adds during each association event. To study telomere elongation *in vivo* we have established a live-cell imaging assay to track individual telomerase RNPs in HeLa cells. Using this assay and the drug imetelstat, which is a competitive inhibitor of telomeric DNA binding, we demonstrate that stable association of telomerase with the single-stranded overhang of the chromosome end requires telomerase-DNA base-pairing. Furthermore, we show that telomerase processivity contributes to telomere elongation *in vivo*. Together, these findings provide new insight into the dynamics of telomerase recruitment and the importance of processivity in maintaining telomere length in human cancer cells.

## Introduction

Chromosomes in human cells are capped by telomeres, repetitive DNA tracts bound by the shelterin protein complex (de Lange, 2005). Telomeres shorten during each cell cycle due to the failure of the DNA-replication machinery to copy the very end of each chromosome (Harley *et al.*, 1990). To counteract this shortening, continuously dividing cells, such as stem cells and most cancer cells, express telomerase (Stewart and Weinberg, 2006; Schmidt and Cech, 2015). Telomerase is an RNA-containing reverse transcriptase, which adds DNA to the 3’ single-stranded overhang of human chromosomes specified by the template region of the telomerase RNA (TR) (Cech, 2004).

The requirement of cancer cells to express telomerase is highlighted by the frequent occurrence of mutations in the promoter of the gene for telomerase reverse transcriptase (TERT) (Horn *et al.*, 2013; Huang *et al.*, 2013). These mutations activate the mono-allelic expression of TERT (Bell *et al.*, 2015; Borah *et al.*, 2015; Stern *et al.*, 2015; Chiba *et al.*, 2017), which is normally down-regulated when human cells differentiate. In addition to its importance in cancer formation and survival, defects in telomerase-mediated telomere maintenance are associated with a number of premature aging diseases, such as Dyskeratosis Congenita (Armanios and Blackburn, 2012). Thus, telomere maintenance by telomerase plays a key role in several human pathologies, and understanding its basic biology could lead to new approaches to treat these diseases.

Human telomerase is a ribonucleoprotein (RNP), composed of the TERT protein and the telomerase RNA (TR). In addition, the telomerase holoenzyme contains accessory subunits, for example dyskerin, NHP2, and NOP10, which associate with TR to stabilize the RNA and ensure its nuclear localization (Schmidt and Cech, 2015). *In vitro*, human telomerase can processively synthesize multiple telomeric repeats without dissociating from its DNA substrate (Wu *et al.*, 2017b). *In vivo*, telomerase is thought to add ~50-60 nucleotides to most chromosome ends in a single processive step (Zhao *et al.*, 2009). In support of this hypothesis, recent results have implicated telomerase processivity as an important contributor to telomere maintenance *in vivo* (Wu *et al.*, 2017a). The nucleus of a human cancer cell only contains ~250 fully assembled telomerase RNPs (Xi and Cech, 2014), which is approximately stoichiometric with the number of chromosome ends after DNA replication has occurred.

Telomerase is recruited to telomeres during the S-phase of the cell cycle by a direct interaction between the OB-fold domain of the shelterin component TPP1 and the telomerase essential N-terminal (TEN)-domain of TERT (Nandakumar and Cech, 2012; Sexton *et al.*, 2012; Zhong *et al.*, 2012; Schmidt *et al.*, 2014). Using live cell single-molecule imaging and telomerase with a 3xFLAG-HaloTag (referred to as Halo or HaloTag throughout the rest of the manuscript), we have recently demonstrated that telomerase rapidly diffuses through the nucleus of human cells, searching for telomeres to bind (Schmidt *et al.*, 2016). When telomerase encounters a chromosome end it can form two types of interactions: short “probing” interactions and long “static” interactions. Importantly, the specific binding of TPP1 and TERT is required for the formation of both types of interactions. We postulated that in addition the long-static interactions require base-pairing of TR to single-stranded telomeric DNA and therefore represent telomerase RNPs that are actively elongating the telomere, but we did not provide direct evidence for this hypothesis.

Here we demonstrate that long-static interactions indeed require base-pairing of TR with the chromosome end by utilizing the cancer drug imetelstat (JNJ-63935937, also known as GRN163L), a first-in-class telomerase inhibitor currently in clinical development in hematologic malignancies. Imetelstat, a 13-mer thiophosphoramidate oligonucleotide, is complementary to the template region of TR and prevents its base-pairing with telomeric DNA (Herbert *et al.*, 2005). Furthermore, we demonstrate that Halo-telomerase, which has normal activity but reduced processivity, elongates telomeres at a lower rate than wild-type telomerase in cells; this highlights the importance of the intrinsic processivity of telomerase for telomere maintenance. Together, these observations provide new insight into telomerase recruitment to telomeres and the contribution of telomerase processivity to telomere maintenance.

## Results

### Halo-telomerase is active but has reduced processivity

Because our live cell single-molecule imaging utilizes Halo-TERT, we determined the enzymatic properties of telomerase modified with N-terminal tags. We overexpressed tagged TERT proteins with TR in HEK293T cells (Fig. 1A). WT TERT and Halo-TERT associated with similar amounts of TR, indicating that the HaloTag does not disrupt the assembly of TERT and TR (Fig. 1B). To measure the catalytic properties of Halo-telomerase, we carried out direct telomerase extension assays (Fig. 1C). While telomerase activity normalized to the number of cells used as input material was increased in Halo-TERT samples relative to WT TERT (Fig. 1D), normalization to the amount of TERT purified showed similar amounts of telomerase activity (Fig. 1E). The presence of a 3xFLAG-tag led to a small decrease in processivity, while the HaloTag led to a larger ~20% reduction, both decreases being statistically significant (Fig. 1F). The Halo-telomerase had the same activity and processivity with or without the fluorescent dye used for live cell imaging. Furthermore, the HaloTag did not affect the functional interaction of telomerase with the telomeric protein TPP1, demonstrated by the increase in processivity in the presence of POT1/TPP1 (Fig. 1G,H). We conclude that the introduction of the HaloTag reduces telomerase processivity, as previously shown for Halo- and 3xFLAG-TERT (Chiba *et al.*, 2016; Schmidt *et al.*, 2016). Importantly, the HaloTag does not affect telomerase activity, RNP assembly, or the interaction of TERT with its telomeric partner, TPP1.

**Figure 1.**
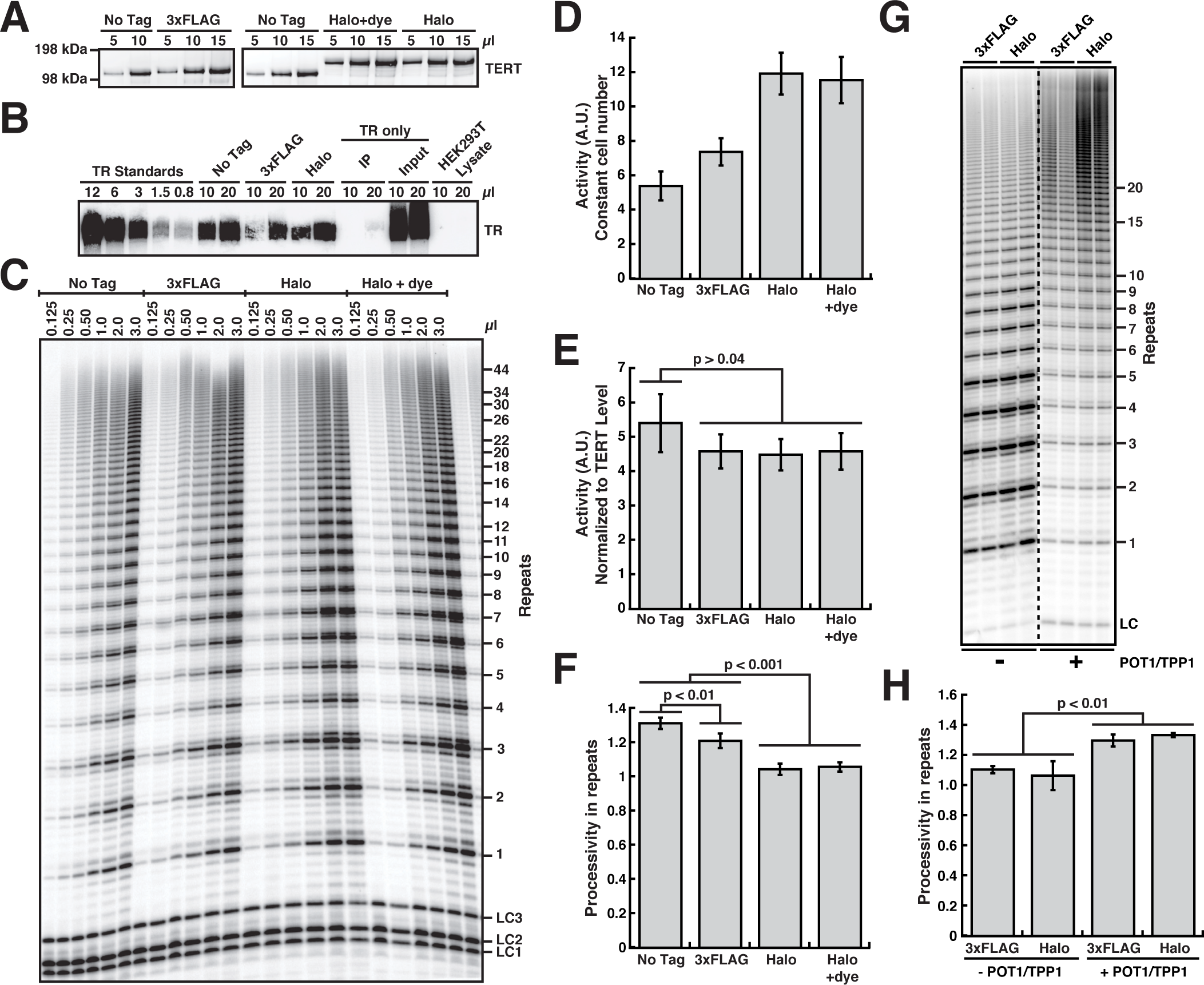
The HaloTag on TERT has no effect on telomerase properties except for a reduction in processivity. (A) Western blot of telomerase immuno-purified from HEK293T cells over-expressing various untagged and tagged TERT proteins and TR, probed with an anti-TERT antibody. (B) Northern blot of RNA extracted from immuno-purified telomerase variants, probed with three TR probes. TR only input and IP were derived from HEK293T cells overexpressing only TR but not TERT, to control for non-specific TR purification. (C) Direct telomerase extension assay at 50 mM KCl of various immuno-purified telomerase variants. LC1, LC2, and LC3, labeled DNA loading controls. (D) Quantification of telomerase activity normalized to the loading controls and the number of cells used as input for immuno-purification (n = 6, mean ± SD). (E) Quantification of telomerase activity normalized to the loading controls and the TERT level (see panel A, n = 6, Mean ± SD, t-test). (F) Quantification of telomerase processivity using the decay method (n = 5, Mean ± SD, t-test). (G) Direct telomerase extension assay at 150 mM KCl (to limit processivity) of 3xFLAG- and 3xFLAG-HaloTag-telomerase immuno-purified from HEK293T cells using anti-FLAG resin in the absence and presence of POT1/TPP1. LC, labeled DNA loading control. (H) Quantification of 3xFLAG- and 3xFLAG-HaloTag-telomerase processivity in the absence and presence of POT1/TPP1 using the decay method (n = 5, Mean ± SD, t-test).

### Halo-telomerase elongates telomeres *in vivo*

To test whether Halo-telomerase can elongate telomeres *in* vivo, we stably introduced WT TERT, Halo-TERT, and Halo-TERT harboring the K78E recruitment-deficient mutation into HeLa cells by retroviral transduction (Fig. 2A). This approach leads to overexpression of the respective *TERT* allele (Fig. 2B), which elicits a dominant effect by outcompeting the endogenous TERT for assembly with TR into the mature telomerase RNP (Fig. 2A). TERT was overexpressed to a similar degree in all polyclonal, virally transduced cell lines (Fig. 2B). To measure the telomerase activity in these cells, we immuno-purified telomerase and subjected it to direct telomerase assays (Fig. 2B, Fig. S1A). Similar to telomerase overexpressed in HEK293T cells (see above), we observed comparable catalytic activity for all TERT variants (Fig. S1B) and a reduction of processivity of telomerase RNPs that were modified with the HaloTag (Fig. S1C). As previously shown (Schmidt *et al.*, 2014), TERT overexpression increased telomerase activity per cell in all cell lines (Fig. S1C). Importantly, Halo-TERT harboring the K78E mutation displayed enzymatic properties that were indistinguishable from its WT counterpart (Fig. 2B, Fig. S1A-C).

**Figure 2.**
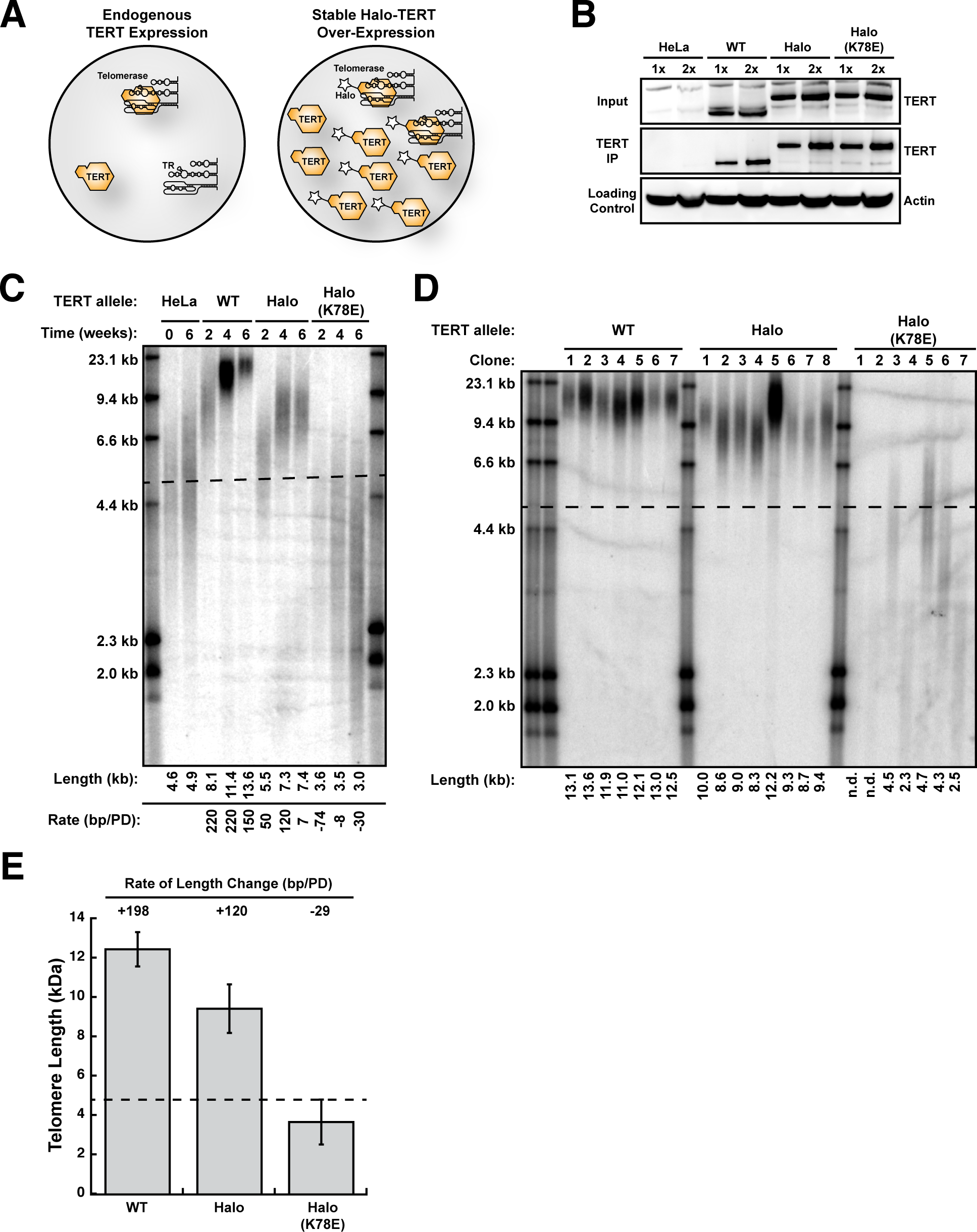
Halo-telomerase elongates telomeres *in vivo*. (A) Experimental design. Over-expression of TERT will increase telomerase levels in HeLa cells by driving the assembly of free TR into telomerase RNPs. Due to its higher levels, exogenous TERT will outcompete endogenous TERT for assembly with TR. (B) Western blots of cell lysates (Input) and TERT immuno-purified (TERT IP) from cell lines over-expressing various *TERT* alleles probed with an anti-TERT antibody. Cell lysates were probed with an anti-beta-Actin antibody as loading control. (C) Telomere length analysis of polyclonal HeLa cell lines stably over-expressing various TERT proteins by Southern Blot of telomeric restriction fragments. Each rate of telomere extension was calculated relative to the previous time point recorded. (D) Telomere length analysis of single cell clones of HeLa cells stably over-expressing various TERT proteins by Southern Blot of telomeric restriction fragments. (E) Quantification of the rate of telomere length change by averaging the telomere length of all single cell clones (see panel D), calculating their length relative to those of the parental HeLa cells (see panel C), and dividing by the number of population doublings between introduction of the *TERT* transgene and analysis of telomere length.

To determine if Halo-telomerase can elongate telomeres in cells, we measured telomere lengths in virally transduced cell lines by Southern blotting. It is important to note that although TERT is substantially overexpressed, the TR subunit becomes limiting so telomerase activity increases only ~1.5-2-fold as seen previously (Cristofari *et al.*, 2007; Schmidt *et al.*, 2014; Xi and Cech, 2014). Telomere length in the parental HeLa cells remained constant over the time course of the experiment (Fig. 2C). Expression of WT TERT led to telomere elongation from ~4.6 kb to ~13.6 kb over the time course of 6 weeks, which corresponds to a growth rate of 150-220 base pairs per population doubling (bp/PD) (Fig. 2C). Telomere length in cells expressing Halo-TERT also increased (from ~4.6 kb to 7.4 kb), but at a slower rate of ~50-120 bp/PD, and telomeres reached their new length set point by 4 weeks (Fig. 2C). Importantly, telomere length in cells expressing Halo-TERT harboring the K78E mutation, which has full enzymatic activity (Fig. S1A-C) but cannot localize to telomeres (Schmidt *et al.*, 2014), shrunk from ~4.6 kb to ~3.0 kb over the six-week time course (Fig. 2C), confirming that TERT overexpressed from the transgene is dominant over endogenous TERT.

As an additional approach to determine the impact of TERT overexpression on telomere length, we isolated single cell clones from the polyclonal populations one week after viral transduction and determined their telomere length 5 weeks after introduction of the *TERT* transgene (Fig. 2D). Telomeres in clones expressing WT TERT and Halo-TERT grew to an average of ~12.5 kb and ~9.4 kb, corresponding to growth rates of ~200 bp/PD and ~120 bp/PD, respectively (Fig. 2E). These growth rates are consistent with those observed in the polyclonal cell populations (Fig. 2C). Clones expressing K78E Halo-TERT shortened to ~3.7 kb at a rate of ~30 bp/PD (Fig. 2D,E). In total, these observations demonstrate that Halo-telomerase elongates telomeres *in vivo*, but it does so at a reduced rate compared to WT telomerase.

### Imetelstat prevents the association of telomerase with its ssDNA substrate

Imetelstat is complementary to the template region of TR and therefore should be a competitive inhibitor of single-stranded telomeric DNA binding to telomerase (Herbert *et al.*, 2005). To test this hypothesis, we established a single-molecule telomerase primer-binding assay (Fig. 3A). Halo-telomerase purified from HEK293T cells was modified with a HaloTag-ligand conjugated to a biotin molecule (Fig. 3B), to allow immobilization on a coverslip surface derivatized with neutradivin (Fig. 3A). Primer binding by telomerase was analyzed by telomerase-dependent recruitment of a fluorescently labeled telomeric oligonucleotide to the surface of the coverslip, visualized by TIRF microscopy (Fig. 3A,C). Importantly, a large fraction (~50%) of the telomerase RNPs immobilized by this approach were enzymatically active, as determined by a single-molecule telomerase activity assay previously established by Sua Myong and colleagues (Fig. 3A,C)(Hwang *et al.*, 2014). We then carried out this single-molecule primer-binding assay in the presence of increasing concentrations of imetelstat (Fig. 3D,E). Imetelstat progressively decreased the amount of telomeric primer bound to telomerase (Fig. 3D,E), consistent with the hypothesis that imetelstat competes with ssDNA for telomerase binding. Half-maximal inhibition (IC_50_) of primer binding occurred at ~16 nM imetelstat (Fig. 3E), which is comparable to the concentration (20 nM) of telomeric primer used in this experiment.

**Figure 3.**
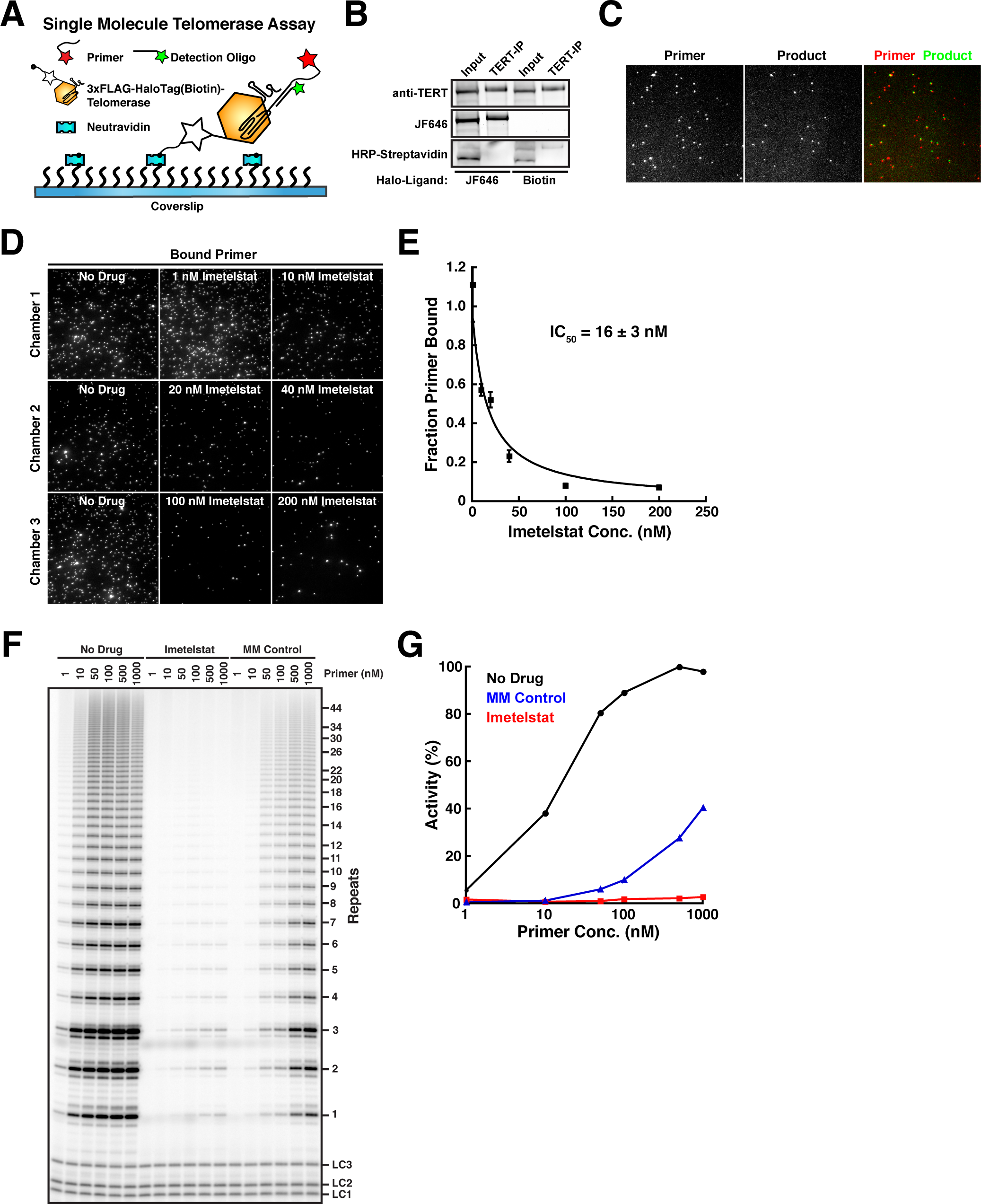
Imetelstat (GRN163L) is a competitive inhibitor of primer-substrate binding by telomerase. (A) Experimental design of single-molecule telomerase primer binding and activity assay. Halo-telomerase is modified with a biotin-HaloTag-ligand and immobilized on the coverslip surface using NeutrAvidin. Primer binding is visualized by telomerase dependent recruitment of a fluorescent primer to the coverslip surface. The telomerase extension product is detected using a fluorescently labeled oligonucleotide anti-sense to the telomerase extension product. (B) Western blot and fluorescence imaging of Halo-telomerase modified with a fluorescent dye (JF646) or biotin, probed with an anti-TERT antibody or HRP-conjugated to streptavidin. (C) Single-molecule TIRF imaging of primer molecules recruited to the coverslip surface by telomerase and its co-localization with telomerase extension products after incubation with nucleotide substrate. (D) Single-molecule TIRF imaging of primer binding by telomerase in the presence of increasing concentrations of imetelstat. (E) Quantification of primer binding to telomerase as a function of imetelstat concentration (5 fields of view per concentration, data points plotted as Mean ± SD, error on IC_50_ reflects error in the corresponding fit of the data to a simple binding curve). (F) Direct telomerase assay at 150 mM KCl in the absence and presence of imetelstat (10 nM), or mismatched control oligonucleotide (MM Control, 10 nM), and increasing concentrations of primer substrate. LC1, LC2, and LC3, labeled DNA loading controls. (G) Quantification of telomerase activity as a function of primer concentration in absence and presence of imetelstat (10 nM), or mismatched control oligonucleotide (MM Control, 10 nM).

As an alternative approach to address how imetelstat affects primer binding to telomerase, we carried out direct telomerase assays in the presence of imetelstat or a control oligonucleotide that carries mismatches at four positions in the nucleotide sequence at varying concentrations of telomeric substrate primer (Fig. 3F)(Asai *et al.*, 2003). Imetelstat (10 nM) strongly inhibited telomerase at all primer concentrations used (Fig. 3F,G). The mismatched control also inhibited telomerase activity, but activity was recovered at higher primer concentrations (Fig. 3F,G), demonstrating that the control compound is a less effective inhibitor of telomerase activity. Together, these results demonstrate that imetelstat acts as competitive inhibitor of telomerase binding to its ssDNA substrate.

### Imetelstat inhibits telomerase *in vivo*

Because we planned to use imetelstat as an inhibitor of telomerase-telomere base-pairing in live cell imaging experiments, we tested how the drug affected telomerase RNP activity *in vivo*. We treated HeLa cells expressing Halo-TERT from the endogenous *TERT* locus with 2 µM of imetelstat or mismatch (MM) control oligonucleotide for 24 h and immuno-purified telomerase RNPs from the treated cells. Similar amounts of TERT and TR were purified from treated and untreated cells (Fig. 4A,B). This demonstrates that a 24 h treatment with imetelstat does not affect telomerase RNP assembly in HeLa cells. While telomerase from untreated cells and cells treated with the mismatch control oligonucleotide showed similar telomerase activity, no activity was detected in telomerase preparations from cells treated with imetelstat (Fig. 4C,D). Together, these observations demonstrate that 24 h of treatment with imetelstat completely inhibits telomerase activity in HeLa cells, without affecting telomerase RNP assembly. In addition, imetelstat likely has a very slow dissociation rate from telomerase, since no telomerase activity is recovered even after a prolonged purification procedure (~3 h).

**Figure 4.**
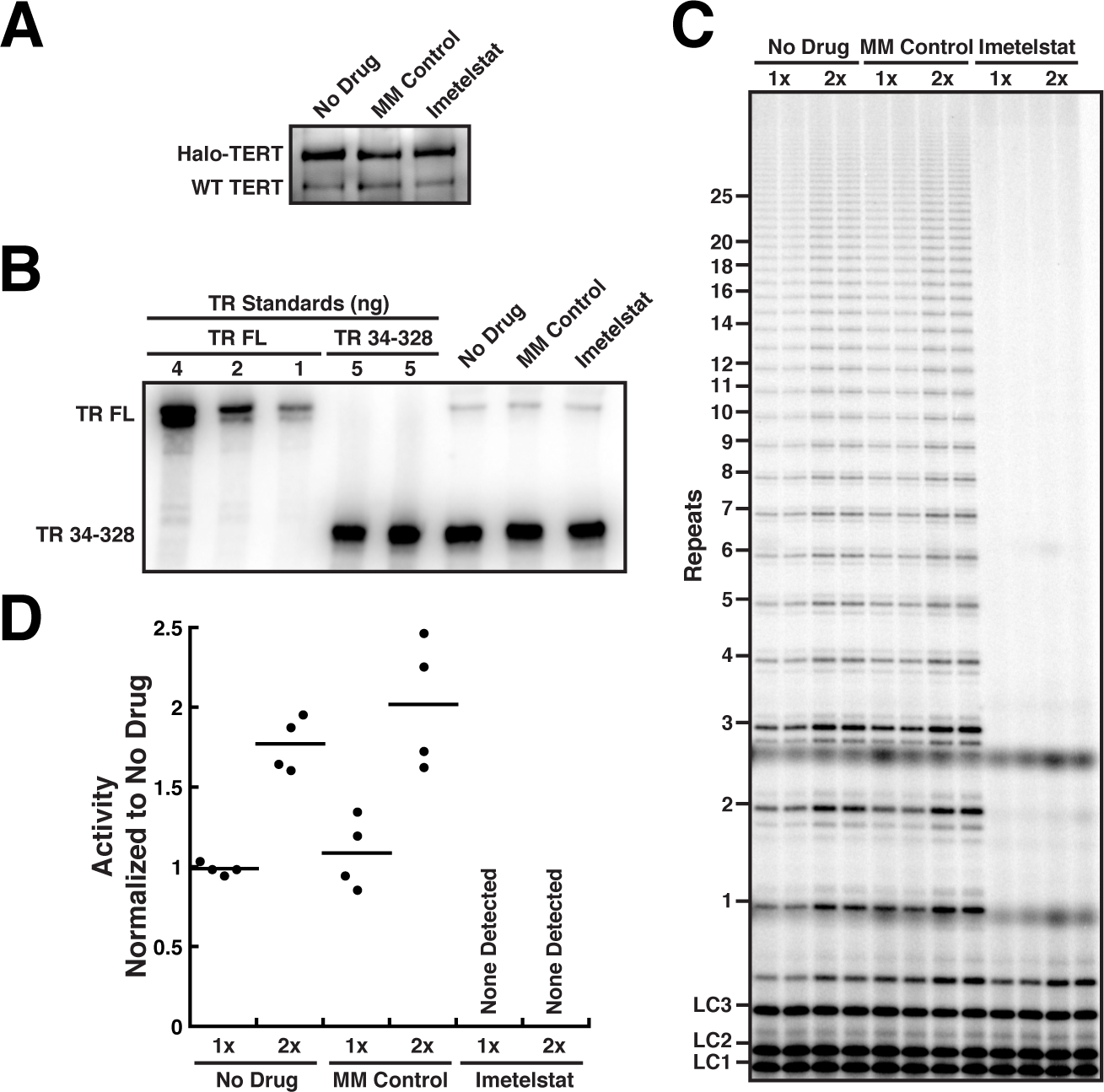
Imetelstat inhibits telomerase *in vivo* but does not affect RNP assembly. (A) Western blot of telomerase immuno-purified from HeLa cells expressing Halo-TERT from the endogenous *TERT* locus, after 24-hour treatment with imetelstat or mismatched control oligonucleotide, probed with an anti-TERT antibody. (B) Northern blot of RNA extracted from telomerase immuno-purified from HeLa cells expressing Halo-TERT after 24-hour treatment with 2 µM imetelstat, 2 µM mismatched control oligonucleotide, or no drug. *In vitro* transcribed full-length (FL) TR was included as size standard and truncated TR 34-328 as loading and recovery control. Blots were probed with three radiolabeled oligonucleotides antisense to TR. (C) Direct telomerase assay at 150 mM KCl of telomerase immuno-purified from HeLa cells expressing Halo-TERT from the endogenous *TERT* locus, after 24-hour treatment with 2 µM imetelstat, 2 µM mismatched control oligonucleotide, or no drug. LC1, LC2, and LC3, labeled DNA loading controls. (D) Quantification of the activity of telomerase purified from HeLa cells expressing Halo-TERT from the endogenous *TERT* locus, after 24-hour treatment with 2 µM imetelstat or 2 µM mismatched control oligonucleotide, normalized to No Drug and loading control (n = 4, Mean).

### Imetelstat inhibits the formation of long-static telomerase–telomere interactions *in vivo*

We previously demonstrated that telomerase forms two types interactions with telomeres: short, dynamic “probing” interactions, and long-lasting static interactions (Fig. 5A) (Schmidt *et al.*, 2016). We speculated that the long-lasting static interactions represent telomerase RNPs that are base-paired with the single-stranded overhang of the chromosome end, but were not able to provide direct evidence for this hypothesis (Schmidt *et al.*, 2016). To test the base-pairing hypothesis, we carried out live cell single-molecule imaging of telomerase trafficking in HeLa cells in the presence of 2 μM imetelstat or the mismatched control oligonucleotide (Fig. 5B, Movies 1-3). We observed dynamic “probing” and long-static interactions under all conditions (Fig. 5C, Movies 1-3), and the overall distribution of diffusion coefficients was unaffected by imetelstat treatment (Fig. 5D), indicating no gross changes in RNP assembly or behavior. However, the number of cells in which we observed long-static interactions was significantly (p < 0.01) reduced in the presence imetelstat in a dose-dependent manner (Fig. 5E,F), but remained unchanged when cells were treated with the mismatched control oligonucleotide (Fig. 5E). These observations demonstrate that the long-static interactions depend on the ability of telomerase to base-pair with the single-stranded overhang of the chromosome end.

**Figure 5.**
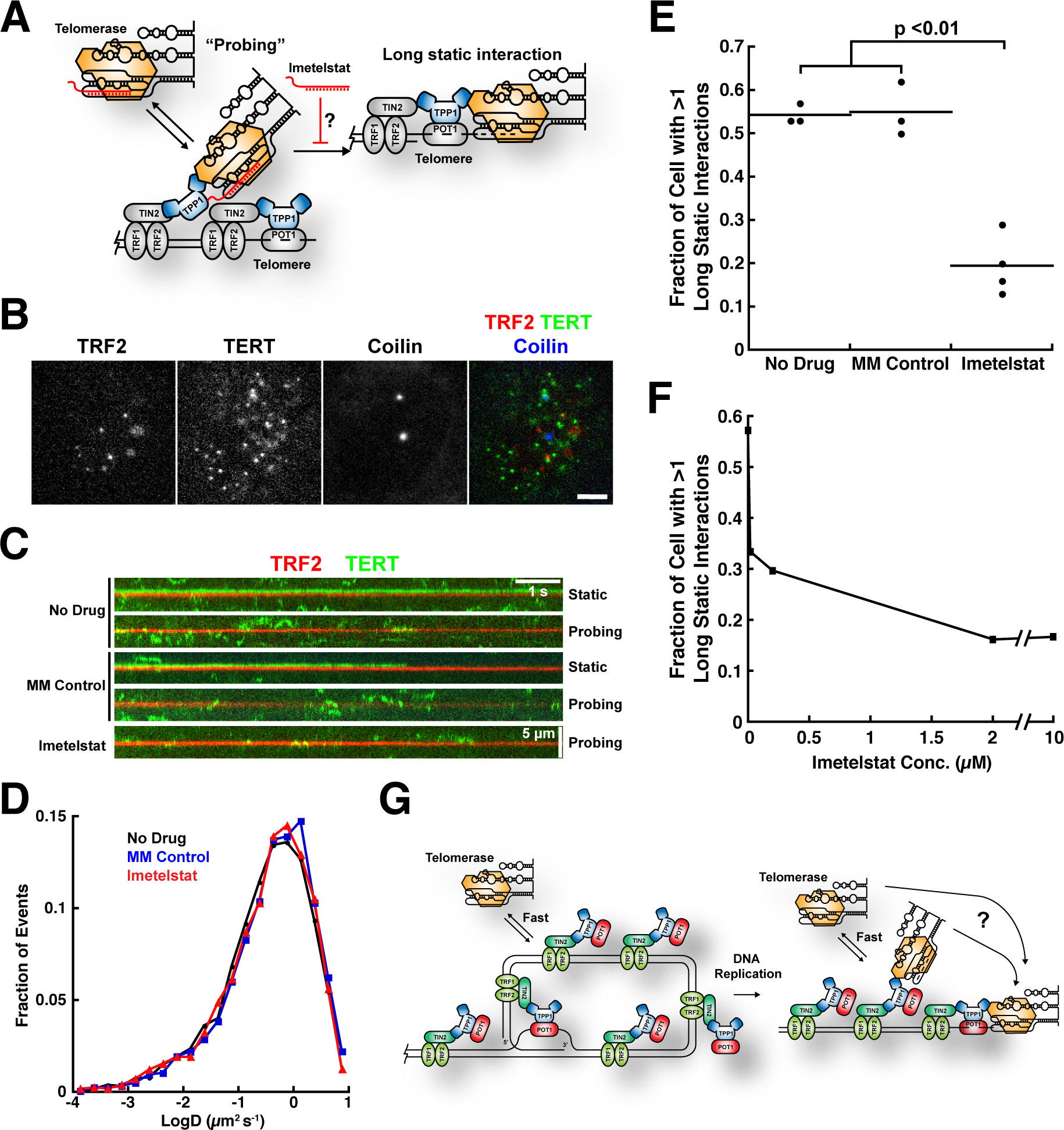
Imetelstat inhibits the formation of long-static telomerase–telomere interactions. (A) Experimental rationale. If long-static telomerase–telomere interactions require the base pairing of TR to the chromosome end, imetelstat should prevent their formation. (B) Still images from three-color live-cell single-molecule imaging experiments. Telomeres were marked by mEOS3.2-TRF2, telomerase was detected using a 3xFLAG-HaloTag-TERT conjugated to JF646, and Cajal bodies were visualized using BFP-coilin (scale bar = 5 µm). (C) Kymographs of TERT and TRF2 trajectories from HeLa cells stably expressing 3xFLAG-HaloTag-TERT and mEOS3.2-TRF2 from their respective endogenous loci. Cells were treated with 2 µM imetelstat, 2 µM mismatched control oligonucleotide, or no drug. (D) Diffusion coefficient distributions of TERT trajectories from HeLa cells stably expressing 3xFLAG-HaloTag-TERT treated with imetelstat, mismatched control oligonucleotide, or no drug (n = 4000-5000 trajectories per condition). (E) Quantification of the fraction of cells with at least one long-static telomerase–telomere interaction after treatment with 2 µM imetelstat, 2 µM mismatched control oligonucleotide, or no drug (N_No Drug_ = 3, N_MM Control_ = 3, N_imetelstat_ = 4, n = 13-36 cells per condition, Mean ± SD, t-test). (F) Quantification of the fraction of cells with at least one long-static telomerase–telomere interaction as a function of imetelstat concentration (n = 28-44 cells per condition). (G) Model for telomerase recruitment to chromosome ends derived from the current study. Short probing interactions require TERT–TPP1 protein-protein binding, while the productive interaction of telomerase with the DNA 3’-end (far right) also requires base-pairing with the template region of TR. Whether telomerase can hop or slide to the 3’ end from internal binding sites or, alternatively, is released back into the nucleus and must collide directly with the 3-terminus is an open question (?).

Another part of our hypothesis was that the short “probing” interactions would not be affected by treating cells with imetelstat, because they are stabilized by protein-protein interactions. We therefore carried out a kinetic analysis of the residence times of TERT particles in proximity to telomeres, Cajal bodies, and other nuclear locations (Fig. S2A-C). All residence time distributions fit well to the sum of two exponential decay functions (Fig. S2A-C), one with a very rapid off-rate and one with a ~10-fold slower off-rate, indicating that two distinct molecular processes underlie the behavior of the TERT RNPs at all nuclear locations. We propose that the fast component reflects TERT particles that are not actually associated with any sub-nuclear structure and by chance did not diffuse very far between two consecutive frames. Consistent with this interpretation, the half-lives of the fast components are similar to the sampling rate of the experiment, ~18-27 ms and 22 ms respectively (Fig. S2E). In contrast, the slow component likely represents TERT particles that are forming an interaction with a sub-nuclear structure. Importantly, the slower off-rate of TERT particles at telomeres, Cajal bodies, and other nuclear locations are similar under all experimental conditions (Fig. S2D), indicating that the underlying molecular interactions are unaffected by treatment with imetelstat. Furthermore, the off-rate of TERT particles at telomeres and Cajal bodies was significantly (p < 0.05) lower than at other nuclear locations (Fig. S2D), consistent with it forming specific interactions with telomeres and Cajal bodies. The fraction of TERT particles that dissociate from telomeres with a slower rate-constant was reduced after treatment with imetelstat (Fig. S2F). This decrease is likely due to the contributions of long-static interactions to this analysis, because these are reduced by imetelstat treatment. In total, these observations demonstrate that the long-static interactions we observe by live cell single-molecule imaging represent telomerase RNPs that are engaged with the telomere by base pairing of TR of the chromosome end.

## Discussion

### Long-Static Telomerase-Telomere Interactions Require Base-Pairing of TR with the Chromosome End

Telomere maintenance is essential for the proliferation of all actively dividing cells in the human body, including stem cells and cancer cells (Stewart and Weinberg, 2006). Telomerase compensates for telomere shrinkage that occurs during semi-conservative DNA replication by adding telomeric repeats to the chromosome ends (Schmidt and Cech, 2015). Telomerase is a unique reverse transcriptase, which synthesizes DNA using the template sequence present in its RNA subunit TR (Wu *et al.*, 2017b). A critical step in telomere lengthening is telomerase recruitment to telomeres, which is not trivial due to the low abundance of telomerase in human cancer cells (Xi and Cech, 2014). We have previously described two different types of interactions that telomerase can from with telomeres (Schmidt *et al.*, 2016): short dynamic “probing” interactions and long-lasting static interactions. Both types of interactions require binding of the TEN-domain of TERT with the TEL-patch of TPP1, because both are eliminated by a point mutation in TERT (K78E).

Imetelstat, a lipid-modified thio-phosphoramidate oligonucleotide that is complementary to the template region of TR (Herbert *et al.*, 2005), allowed us to analyze the contributions of this region of TR to the interactions that we observe *in vivo*. We proposed that the long-static TERT-telomere associations represent telomerase RNPs that are base-paired with the chromosome end and are actively elongating the telomere. Since imetelstat is a competitive inhibitor of the association of telomerase with its DNA substrate, it should interfere with the formation of long-static interactions if they required base-pairing of TR with the chromosome end. Indeed, the frequency of the formation of long-static interactions was dramatically reduced in the presence of imetelstat but remained unchanged when cells were treated with a mismatched control oligonucleotide. These observations support the hypothesis that RNA-DNA base-pairing between TR and the chromosome end is necessary the formation of long-static interactions. Therefore, since telomerase RNPs that are engaged in long-static interactions are base-paired to the chromosome end, they are likely actively elongating the telomere.

### Telomerase Processivity Contributes to Telomere Maintenance *In Vivo*

To processively synthesize multiple telomeric repeats, the template region of TR must be repositioned relative to the substrate DNA in a step called translocation. *In vitro* human telomerase can processively synthesize multiple telomeric repeats, but how much processivity contributes to telomere maintenance *in vivo* remains unclear (Wu *et al.*, 2017b). *In vivo*, the amount of telomere lengthening that occurs at a given telomere depends on two key variables: the number of times that telomerase binds to the DNA at the chromosome end per cell cycle (frequency), and the number of repeats telomerase adds to the telomere per association event (processivity).

The introduction of the HaloTag on the N-terminus of TERT allowed us to visualize telomerase in living cells and inadvertently generated a TERT variant with reduced telomerase processivity without affecting activity, which is the number of nucleotides added per unit time. We could therefore test the contribution of telomerase processivity to telomere elongation *in vivo* by comparing Halo-telomerase with the wild-type RNP. The presence of the HaloTag led to a substantial reduction in the rate of telomere lengthening and in the plateau length when TERT was overexpressed in human cancer cells. Overexpression of telomerase likely leads to telomere growth by increasing the frequency of lengthening events that occur per telomere in a given S-phase. The intrinsic processivity of telomerase dictates how many repeats are added in each individual telomere lengthening event, and is unlikely to be affected by overexpressing TERT. Under the conditions in our experiments, the amount of TERT protein vastly exceeds the amount of TR, making TR the limiting component for telomerase assembly. Therefore, the telomerase RNP concentrations and thus the frequency of telomere lengthening events are likely similar for all *TERT* alleles tested. The difference in telomere lengthening can therefore be attributed to the difference in intrinsic processivity of WT *versus* Halo-TERT. Importantly, telomere lengthening depended on the ability of Halo-TERT to localize the telomeres, demonstrating that even though it occurs at a lower overall rate, telomere elongation is carried out by Halo-telomerase. We conclude that a small decrease in telomerase processivity (~20 %) can have a substantial effect on telomere length when aggregated over multiple cell divisions. This observation highlights the importance of telomerase processivity for telomere lengthening *in vivo*, consistent with recent observations made by others (Wu *et al.*, 2017a).

### Implications for Telomere Maintenance

We have now defined elements of the molecular basis of the short “probing” and long-static interactions observed in our live cell single-molecule imaging experiments. Because the protein-protein interaction between TERT and TPP1 is required for both types of interactions and base-pairing of TR with the chromosome end only for the formation of long-static interactions, it is tempting to speculate that “probing” interactions are an intermediate between free telomerase and telomerase that is elongating the telomere (Fig. 5G). Probing the telomere would then increase the chance of telomerase finding the chromosome end, and because it is transient it would not trap the small pool of telomerase RNPs at telomeres that do not have the 3' overhang available for binding. If telomere elongation were only possible during a brief window of time after DNA replication (Zhao *et al.*, 2009), this “probing” mechanism would increase the probability of telomerase finding the chromosome end during this time window (Fig. 5G). Whether telomerase is recruited to specific telomeres (e.g., short telomeres) during a particular time window will be the subject of future investigation. For now, the results presented in this study lay the groundwork for a comprehensive quantitative analysis of telomere elongation by telomerase in human cancer cells.

## Materials and Methods

### Plasmids Construction

Plasmids for the expression of WT- and 3xFLAG-TERT were previously described. The plasmid for overexpression of 3xFLAG-HaloTag TERT was generated by first ligating *TERT* (amplified with TERT for and TERT rev, see Table 1) into the pHTN HaloTag CMV-neo vector (Promega) cut with EcoRI and NotI (NEB). The sequence coding for the 3xFLAG-HaloTag was amplified from genomic DNA of a HeLa cell line stably expressing 3xFLAG-HaloTag-TERT from the endogenous *TERT* locus (amplified with 3xFLAG-HaloTag for and 3xFLAG-HaloTag rev, see Table 1), and ligated into pHTN containing *TERT* using NheI and EcoRI. To generate the exact coding sequence present in the HeLa cell line stably expressing 3xFLAG-HaloTag-TERT from the endogenous *TERT* locus, the EcoRI site was mutated to a KpnI site using quick change (amplified with EcoRI QC for and EcoRI QC rev, see Table 1). Plasmids for generation of retroviruses coding for various *TERT* alleles were generated by Gibson assembly, inserting a *TERT* (amplified with TERT Gibson for and TERT Gibson rev, see Table 1) or *3xFLAG-HaloTag-TERT* (amplified with Halo TERT Gibson for and TERT Gibson rev, see Table 1) fragment into the pBABE PuroR vector (amplified with pBABE Gibson for and pBABE Gibson rev, see Table 1).

**Table 1.**
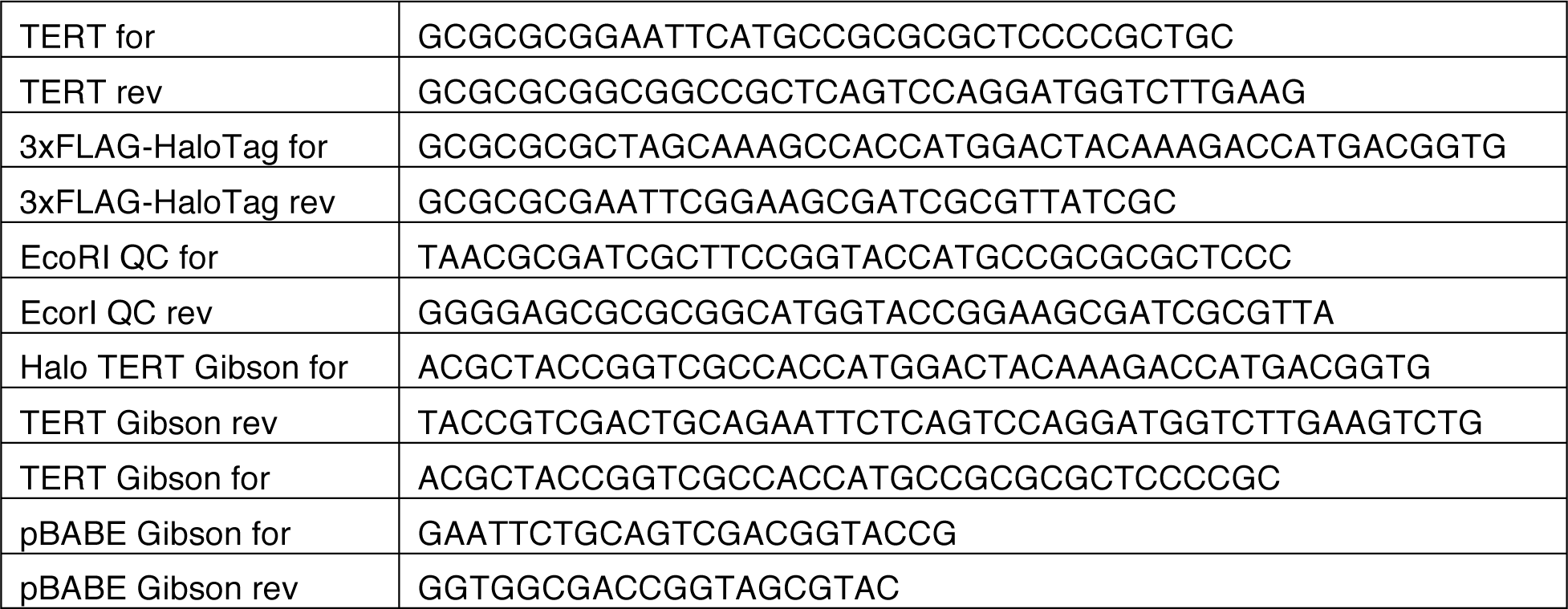
Primers used in this study.

### Cell Culture and Generation of Stable Cell Lines

All cell lines were derivatives of Hela-EM2-11ht and were grown in high glucose Dulbecco’s Modified Eagle Medium (DMEM) supplemented with 10% fetal bovine serum (FBS), 2 mM GlutaMAX^TM^-I (Life Technologies), 100 units/ml penicillin and 100 μg/ml streptomycin at 37 °C with 5% CO_2_. Imaging experiments were carried out in CO_2_-independent media supplemented with 10% FBS, 2 mM GlutaMAX^TM^-I (Life Technologies), 100 units/ml penicillin and 100 μg/ml streptomycin in a humidified imaging chamber heated to 37 °C. For S-phase synchronization, cells were arrested in growth medium containing 2 mM thymidine for 16 h, released for 9 h, followed by a second thymidine 16 h arrest prior to release into S-phase. Puromycin selection was carried out at a concentration of 1 µg/ml (Sigma). Stable cell lines were generated by retroviral transduction as previously described (Schmidt *et al.*, 2014). Single cell clones were derived by diluting a suspension of the polyclonal cells to ~ 6 cells/ml and plating 150 µl of this suspension into each well of a 96 well plate. Only clones that clearly formed a single colony of the appropriate size were used. Both polyclonal and clonal stable cell lines were continuously cultured in media containing puromycin.

Imetelstat and the mismatched control oligonucleotide were a gift of Janssen Research & Development, LLC (920 Route 202, Raritan, NJ 08869). Imetelstat and the mismatched control oligonucleotide were dissolved in PBS at concentrations between 1-2 mM and stored at −20 ºC. Concentrations were verified using OD_260_. HeLa cells were incubated with 2 µM of imetelstat or the mismatched control oligonucleotide for 24 hours prior to imaging or telomerase purification.

Transient transfections of BFP-coilin were carried out with the Nucleofector^TM^ 2b device, using Kit R and the high efficiency protocol for HeLa cells (Lonza).

### Telomerase Purification and Activity Assay

Telomerase was over-expressed as previously described (Sauerwald *et al.*, 2013). FLAG IP was performed with Anti-FLAG® M2 Affinity Gel (Sigma-Aldrich, A2220) using HEK293T cell lysates prepared with CHAPS lysis buffer (10 mM Tris-HCl pH 7.5, 1 mM MgCl_2_, 1 mM EGTA, 0.5% CHAPS, 10% glycerol, 1 mM PMSF, 1 mM DTT).

Telomerase IP of endogenous and over-expressed TERT was carried out with a sheep polyclonal anti-TERT antibody, which was a gift from Scott Cohen (Children’s Medical Research Institute and University of Sydney, Westmead, Australia), from CHAPS lysates of ~100x10^6^ HeLa or HEK293T cells. The HaloTag was labeled during the incubation with the resin using a concentration 0.5 µM JF646 (a kind gift from Luke Lavis, HHMI Janelia Research Campus) or PEG-Biotin (Promega) HaloTag-ligand. The telomerase purifications and activity assays were carried out as previously described (Zaug *et al.*, 2013), using indicated salt (KCl) and substrate concentrations. Purification of POT1/TPP1 and telomerase assays in the presence of POT1/TPP1 were carried out as previously described (Schmidt *et al.*, 2014), using indicated salt (KCl), substrate, and POT1/TPP1 concentrations. Loading controls were phosphorylated telomeric DNA 18- and 21-mers. To analyze the effect of imetelstat on telomerase activity, the activity assay was initiated by adding primer, imetelstat, and nucleotides simultaneously. Telomerase activity was quantified by the total amount of radioactive counts incorporated into products and normalized to the sum of the loading controls. Telomerase processivity was determined by the decay method previously described (Latrick and Cech, 2010). The radioactive counts of each major telomeric repeat product were divided by the number of dG nucleotides incorporated during its synthesis (1 for the first product, 4 for the second product, 7 for the third product, etc.), which converts the radioactive signal into an estimate of the number of molecules present in each band. The number of molecules in each band represents products that have dissociated from telomerase after each round of repeat synthesis. Telomerase processivity was determined by fitting this decay from repeats (2-15), which contain greater than 95% of the measured molecules, excluding the first repeat, to a single exponential decay function Y = A*e^(-k*Repeat)^. Processivity was calculated as ln(2)/k and has units of telomeric repeats.

### Western Blotting

The protein samples were separated on 4-12% Bis-Tris gels (Life Technologies), followed by standard western blotting procedures. TERT was detected by a primary antibody anti-TERT (Rockland Immunochemicals, 600-401-252, 1:1000) and a secondary antibody peroxidase-AffiniPure donkey anti-rabbit IgG (H+L) (Jackson, 711-035-152, 1:2000). The HaloTag modified with biotin was detected using Strepavidin-HRP (Pierce, 1:2000). SuperSignal® West Femto Chemiluminescent Substrate (Thermo Scientific) was used to generate enhanced chemiluminescence signal, which was detected with a FluorChem HD2 imaging system (Alpha Innotech). JF646 fluorescence was detected directly on the acrylamide gel, prior to western blotting, using a Typhoon Trio PhosphorImager (GE Healthcare).

### RNA Extraction and Northern Blotting

To determine the level of TR contained in telomerase immuno-purifications, telomerase elutions (~ 50 µl) were subjected to Trizol (Invitrogen, 0.5 ml) extraction following the manufacturer’s instructions. In some instances, a loading and recovery control TR 34-328 (5 ng per sample) was included in the Trizol reagent. Precipitated RNA was resuspended in 50 µl of formamide loading buffer and half of the sample was separated in a 6% TBE Urea polyacrylamide gel (Life Technologies). RNA was transferred onto a Hybond N+ membrane (GE Healthcare) using a wet-blotting apparatus in 1x TBE for 1 h at 1 amp of current. Following blotting, the RNA was UV-crosslinked to the membrane and incubated in Church buffer for 2 h at 50°C. TR was detected using three DNA oligos (CTTTTCCGCCCGCTGAAAGTCAGCGAG, CTCCAGGCGGGGTTCGGGGGCTGGGCAG, and CGTGCACCCAGGACTCGGCTCACACATG) which were radioactively labeled using T4 polynucleotide kinase (NEB). Probe (10x10^6^ cpm) in Church buffer was incubated with the membrane for at least 2 h. The membranes were washed three times with 2xSSC, 0.1% SDS before exposure to a phosphor-imager screen over-night. Detection was carried out using a Typhoon Trio PhosphorImager (GE Healthcare).

### Telomere Length by Southern Blotting

Telomere restriction fragment length analysis was carried out as previously described (Nandakumar and Cech, 2012; Schmidt *et al.*, 2014), using 3 µg of genomic DNA prepared from HeLa cell lines using the GenElute™ Mammalian Genomic DNA Miniprep kit (Sigma-Aldrich). The rate of telomere length change was calculated by dividing the change in mean telomere length by the number of population doublings (PD), assuming 1.1 PD per day or 21.8 h per PD for the HeLa cells used in these experiments.

### Single Molecule Primer Binding and Telomerase Activity Assay

Surface-passivated Nexterion coverslips (22x22 mm, 170 +/- 5 µm thickness, Schott) were prepared as follows. Coverslips were first cleaned with 3% Alconox (Alconox, Inc.) which was brought to a boil in the microwave for 30 min in a sonicating water bath. After three washes with ddH_2_O, the coverslips were treated with Piranha solution (3 parts conc. H_2_SO_4_, 1 part H_2_O_2_) followed by three ddH_2_O washes. The glass surface was activated in two steps. First the coverslips were sonicated in 1 M fresh KOH for 30 min; after three ddH_2_O washes, the coverslips were dried and subjected to 60 min of UZ/Ozone cleaning (Novascan Technologies, Inc.). Coverslips were silanized by vapor deposition of N-(2-Aminoethyl)-3-Aminopropyl-Triethoxysilane (Gelest, Inc.) in a desiccator for 4-16 h. After silanization the coverslips were derivatized using PEG 5000 – succinimidyl valerate (8% solution in fresh 0.1 M sodium bicarbonate) containing a small amount (~3%) of Biotin-PEG 5000 - succinimidyl valerate (Laysan Bio, Inc.) for 24 h. Imaging chambers were assembled by taping a 18x18 mm coverslip onto a 22x22 mm coverslip using double-sided stick tape and mounted on the microscope using a custom-made holder as previously described (Schmidt *et al.*, 2012). Each coverslip yielded three channels with a volume of 5-10 µl. To prepare the chambers for telomerase immobilization the channels were incubated with NeutrAvidin (50 ng/ml in PBS, Thermo Scientific) for 5 min. Channels were then washed with 5 channel volumes of imaging buffer (50 mM Tris pH 8.0, 50 mM KCl, 1 mM MgCl_2_, 0.5 mg/ml BSA, 0.05% TWEEN-20, 2 mM TROLOX, 0.2 mg/ml glucose oxidase, 0.035 mg/ml catalase, 4.5 mg/ml glucose) and incubated in imaging buffer for 5 min. To analyze primer binding to telomerase, telomerase purified from HEK293T cells expressing 3xFLAG-HaloTag-TERT derivatized with Biotin-PEG-HaloTag-ligand was diluted 1:20 in imaging buffer and incubated with 20 nM of fluorescent primer A5 (Cy3-TTTTTAGGGTTAGCGTTAGGG, IDT) for 5 min. When analyzing the impact of imetelstat on primer binding, primer substrate and imetelstat were added simultaneously to telomerase. The telomerase solution was then loaded into the imaging channels and immobilized for 5 min before washing the channel with 5 channel volumes of imaging buffer. To visualize telomerase activity, immobilized telomerase-primer complexes were incubated with 500 µM each dATP, dTTP, and dGTP, and 10 nM of detection oligonucleotide (Cy5-CCCTAACCCTAACCC, IDT) in imaging buffer for 5 min prior to imaging. TIRF imaging was carried out using Nikon N-STORM microscope equipped with a TIRF illuminator, 405 nm (20 mW), 488 nm (50 mW), 561 nm (50 mW), and 647 nm (125 mW) laser lines, an environmental chamber to control humidity and temperature, two iXon Ultra 897 EMCCD cameras (Andor), a 100x oil-immersion objective (Nikon, NA = 1.49), two filter wheels, and the appropriate filter sets. To analyze primer binding, 10 frames of a given field of view were acquired at 20 frames per second. Average intensity projections of these short image sequences were analyzed using in-house MATLAB code, implementing particle detection code publicly available (https://site.physics.georgetown.edu/matlab/code.html). To determine the number of primer molecules in a given field of view, the number of particles with intensities corresponding to a single fluorophore was determined by fitting the intensity profiles of detected particles to a normal distribution and counting the number of particles within one standard deviation from the mean of this distribution. To determine the fraction of primer molecules bound in the presence of imetelstat, the average particle number of 5 fields of view was divided by the number of particles detected in the absence of drug. One channel of each coverslip used for this experiment was a no drug control to account for coverslip surface variability. To determine the IC_50_ for inhibition of primer binding by imetelstat, the fraction of primer bound was plotted as a function of Imtelstat concentration and fit to a binding curve Fraction bound = 1 – [Drug] / ([Drug] + IC_50_). To image telomerase product formation, primer substrate and product detection oligonucleotide were imaged simultaneously for 10 frames at 20 frames per second. The percentage of active telomerase RNPs was determined as the fraction of primer signals that co-localized with a product signal divided by the total number of primers detected.

### Single-molecule Live Cell Imaging

Three-color single-molecule live cell imaging was carried out as previously described (Schmidt *et al.*, 2016), using the HeLa cell line stably expressing 3xFLAG-HaloTag TERT and mEOS3.2-TRF2 from their respective endogenous loci. Briefly, BFP-coilin was transfected into cells 48 h prior to imaging, followed by a double thymidine block. Imetelstat was added to cells 24 h before imaging, simultaneously with the release from the first thymidine block. 3-4 h after release into S-phase, FLAG-HaloTag-TERT was labeled by subjecting cells to a 2 min pulse of 100 nM JF646 HaloTag-ligand (a kind gift from Luke Lavis) in tissue culture medium (Grimm *et al.*, 2015). BFP-coilin was imaged first for ~1 s under continuous illumination. 3xFLAG-HaloTag-TERT and mEOS3.2-TRF2 (red state) were imaged simultaneously. Movies were acquired for 15 s on a Nikon N-STORM microscope under highly inclined and laminated optical sheet (HILO) conditions (Tokunaga *et al.*, 2008), with a 1.49 NA 100x oil-immersion TIRF objective (Nikon) at 46 frames per second. The two imaging channels were projected onto two iXon Ultra 897 EMCCD cameras (Andor) using TwinCam dual emission image splitter (Cairn). The channels were aligned prior to every imaging session using TetraSpeck^TM^ microspheres (ThermoFisher).

### Single Particle Tracking

Single particle trajectories were generated with MatLab 2011b (Mathworks Inc., USA) using SLIMfast, which implements the Multiple-Target-Tracing algorithm (Sergé *et al.*, 2008; Liu *et al.*, 2014), and evaluated using the script evalSPT (Normanno *et al.*, 2015). Particle detection was carried out using 9x9 pixel detection, error rate of 10^-6^, and one deflation loop. Particle tracking for determining diffusion coefficients was carried out by setting the upper bound of the expected diffusion coefficient to D = 5 µm^2^/s. To determine the life time of short “probing” interactions, the maximal expected diffusion coefficient was set to D = 0.1 µm^2^/s, the maximal OFF-Time was set to 3 frames and intensity fluctuation weight to 0.5. To analyze the binding properties of TERT particles at different nuclear locations, TERT tracks were assigned to telomeres, Cajal bodies, or other nuclear locations as previously described (Schmidt *et al.*, 2016). To determine the dissociation rate of TERT interactions at these locations, the survival probabilities of TERT particles were fit to a double exponential decay function Y = A*e^(-kfast*t)^ + B*e^(-kslow*t)^ using tracks ranging from 0.044 – 1 s (2-46 frames), which encompassed >95% of the detected trajectories. The fraction of the particles that dissociate with the slower rate constant was calculated as Fraction_slow_ = B/(A+B). Long, static interactions were identified by manual inspection of movies from HeLa cells expressing 3xFLAG-HaloTag and mEOS3.2-TRF2, as previously described (Schmidt *et al.*, 2016). All analysis was carried out blinded to whether the particular movie was generated from cells treated with imetelstat, mismatched control oligonucleotide, or no drug.

## Code Availability

All Matlab scripts are available upon request from the authors.

## Acknowledgements

We thank Cech lab members Dan Youmans, Ci Ji Lim, and Yicheng Long for assistance with experiments and useful discussions, Joe Dragavon and the BioFrontiers Advanced Light Microscopy Core for assistance with imaging, and Luke Lavis and Zhe Liu (HHMI Janelia Research Campus) for providing HaloTag dyes and assistance with data analysis. We especially thank Janssen Research and Development, LLC, for the generous gift of imetelstat and the mismatched control oligonucleotide, which were essential for this study. This study was supported by grants to J.C.S (K99GM120386) and T.R.C. (R01GM099705). T.R.C. is an investigator of the Howard Hughes Medical Institute.

## Conflict of interest statement

T.R.C. is on the board of directors of Merck, Inc., which provides no funding for his research.

